# Inherited defects in natural killer cells shape tumor immune microenvironment, clinical outcome and immunotherapy response

**DOI:** 10.1101/471912

**Authors:** Xue Xu, Jianqiang Li, Jinfeng Zou, Xiaowen Feng, Chao Zhang, Ruiqing Zheng, Weixiang Duanmu, Arnab Saha-Mandal, Zhong Ming, Edwin Wang

**Affiliations:** School of Computing, Shenzhen University, Shenzhen, China; University of Calgary, Cumming School of Medicine, Calgary, Alberta, Canada; Princess Margaret Cancer Center, University Health Network, Toronto, Ontario, Canada; Department of Mathematics, Dalian Institute of Technology, Dalian, China; Department of Medicine, McGill University, Montreal, Canada

## Abstract

Tumor immune microenvironment (TIME) plays an important role in metastasis and immunotherapy. However, it has been not much known how to classify TIMEs and how TIMEs are genetically regulated. Here we showed that tumors were classified into TIME-rich, -intermediate and -poor subtypes which had significant differences in clinical outcomes, abundances of tumor-infiltrating lymphocytes (TILs), the degree of key immune programs’ activation, and immunotherapy response across 13 common cancer types (n= ∼6,000). Furthermore, TIME-intermediate/-poor patients had significantly more inherited genetic defects (i.e., functional germline variants) in natural killer (NK) cells, antigen processing and presentation (APP) and Wnt signaling pathways than TIME-rich patients, and so did cancer patients than non-cancer individuals (n=4,500). These results suggested that individuals who had more inherited defects in NK cells, APP and Wnt pathways had a higher risk of developing cancers. Moreover, in the 13 common cancers the number of inheritably defected genes of NK cells was significantly negative-correlated with patients’ survival, TILs’ abundance in TIMEs and immunotherapy response, suggesting that inherited defects in NK cells alone were sufficient to shape TILs’ recruitment, clinical outcome, and immunotherapy response, highlighting that NK cell activation was required in the 13 cancer types to drive the recruitment of immune troops into TIMEs. Thus, we proposed that cancer was a disease of NK cell inherited deficiencies. These results had implications in identifying of high-risk individuals based on germline genomes, implementing precision cancer prevention by adoptive transfer of healthy NK cells, and improving existing immunotherapies by combining of adoptive NK cell transfer (i.e., converting TIME-intermediate/-poor tumors into TIME-rich tumors) and anti-PD-1 or CAR-T therapy.

**Contact:** EW

## Introduction

In the past two decades, classification of tumors based on omic data has resulted in distinct tumor molecular subtypes for each cancer type and then provided a framework to study the molecular mechanisms such as oncogenic pathways and discover drug targets for tumors. Moreover, tumor molecular subtypes enable to inform clinical outcomes and treatments. For example, treatment options can be made based on either Her2+, luminal or basal subtypes of breast cancer (^1, 2, 3, 4^). However, the enormous complexity of cancer subtypes, tumor microenvironment, subclones, and somatic genomic alteration landscapes has been reported. For example, tumor molecular subtypes and patient stratifications based on tumor gene expression profiles showed that each cancer type has its own subtypes which were often unique and not commonly shared between any 2 cancer types. Each subtype has its own oncogenic pathways and somatic mutating drivers.

Tumor microenvironments often interact with tumor cells to affect metastasis and clinical outcomes. In the past few years, immune-checkpoint therapy (ICT) has been able to successfully eliminate tumors in 10-40% of patients with melanoma and other cancer types, however, in the majority of patients, ICT failed to have their intended effect (^5, 6^). It has been believed that understanding of tumor infiltrated immune cells (TILs) in tumor immune microenvironment (TIME) could help in getting insights into ICT response, resistance and might improve existing immunotherapies (^7, 8^). Infiltrating T cells are a critical component for ICT response. However, recently it has been shown that NK (natural killer) cell activation is required in melanoma to recruit CD103+ DC (dendritic cell) and then CD8+ effector T cells. Except for the TIL-T cell recruitment function, the best-known role of NK cells is cancer cell killing and tumor immunosurveillance. To fulfill this function, distinct from T and B cells, NK cell is not mediated by antigen specificity but through multiple germline-encoded activating and inhibiting receptors (^9, 10, 11^), the complex balance of inhibitory and activating signals promotes self-tolerance or drives potent effector function of NK cells.

Historically, tumor molecular subtypes have generated lots of insights into the underlying molecular mechanisms of tumorigenesis, metastasis and informing of treatments. With advances of cancer ICT, CAR-T and other immunotherapies, it has a strong interest in stratifying TIMEs into TIME subtypes. Further stratification of patients on the basis of their TIMEs could discover better insight into overall survival and underlying molecular mechanisms for ICT response and help identify new immunotherapeutic targets. However, so far, we have no idea how to study TIMEs and lack of efficient tools to classify TIMEs. Here we conducted an analysis of the omic data (i.e., tumor RNA-seq and whole-exome sequences of germline genomes) of TCGA (The Cancer Genome Atlas) cancer patients (n=∼6,000) representing 13 common cancer types, and a non-cancer cohort (n=4,500) to show that tumors were classified into three universally distinct TIME subtypes across the 13 common cancer types. They were different in prognosis, TILs’ abundance and degree of immune programs’ activation, regardless of cancer type. Inherited defects of NK cells, antigen processing and presentation (APP) and Wnt pathways in patients’ germline genomes modulated TIME subtypes and ICT response. Importantly, we showed that inherited defects in NK cells alone were sufficient to regulate TILs’ abundance in TIMEs, clinical outcomes and immunotherapy response. Further, individuals who have inherited defects in NK cells, APP and Wnt pathways bear high-risk of developing cancers.

## Results

### Three universal TIME subtypes across 13 common cancers

To classify tumors into TIME subtypes, we applied the unsupervised clustering of gene expression data (i.e., melanoma data from TCGA) based on a set of genome-wide CRISPR–Cas9 screen-determined essential genes (i.e., ICT essential genes, n=1,294) from a previous study (^12^) and other known tumor immune-related genes (^13, 14^) (see Methods). We found that melanoma tumors were classified into three TIME subtypes (Fig 1a). Similarly, the three TIME subtypes were repeatedly obtained in the 13 common cancer types (Supplementary Fig 1): bladder urothelial carcinoma (BLCA), breast invasive carcinoma (BRCA), cervical squamous cell carcinoma and endocervical adenocarcinoma (CESC), colon adenocarcinoma (COAD), head and neck squamous cell carcinoma (HNSC), kidney renal clear cell carcinoma (KIRC), lower grade glioma (LGG), lung adenocarcinoma (LUAD), lung squamous cell carcinoma (LUSC), pancreatic adenocarcinoma (PRAD), skin cutaneous Melanoma (SKCM), stomach adenocarcinoma (STAD), thyroid carcinoma (THCA) and uterine corpus endometrial carcinoma (UCEC). These results suggested that TIME subtypes were much simpler than tumor molecular subtypes, each of which was unique and couldn’t be shared by any two cancer types. Thus, the universal TIME subtypes provided a means to identify subtypes’ unifying features and to understand the underlying common molecular mechanisms of each TIME subtype.

**Figure 1.**
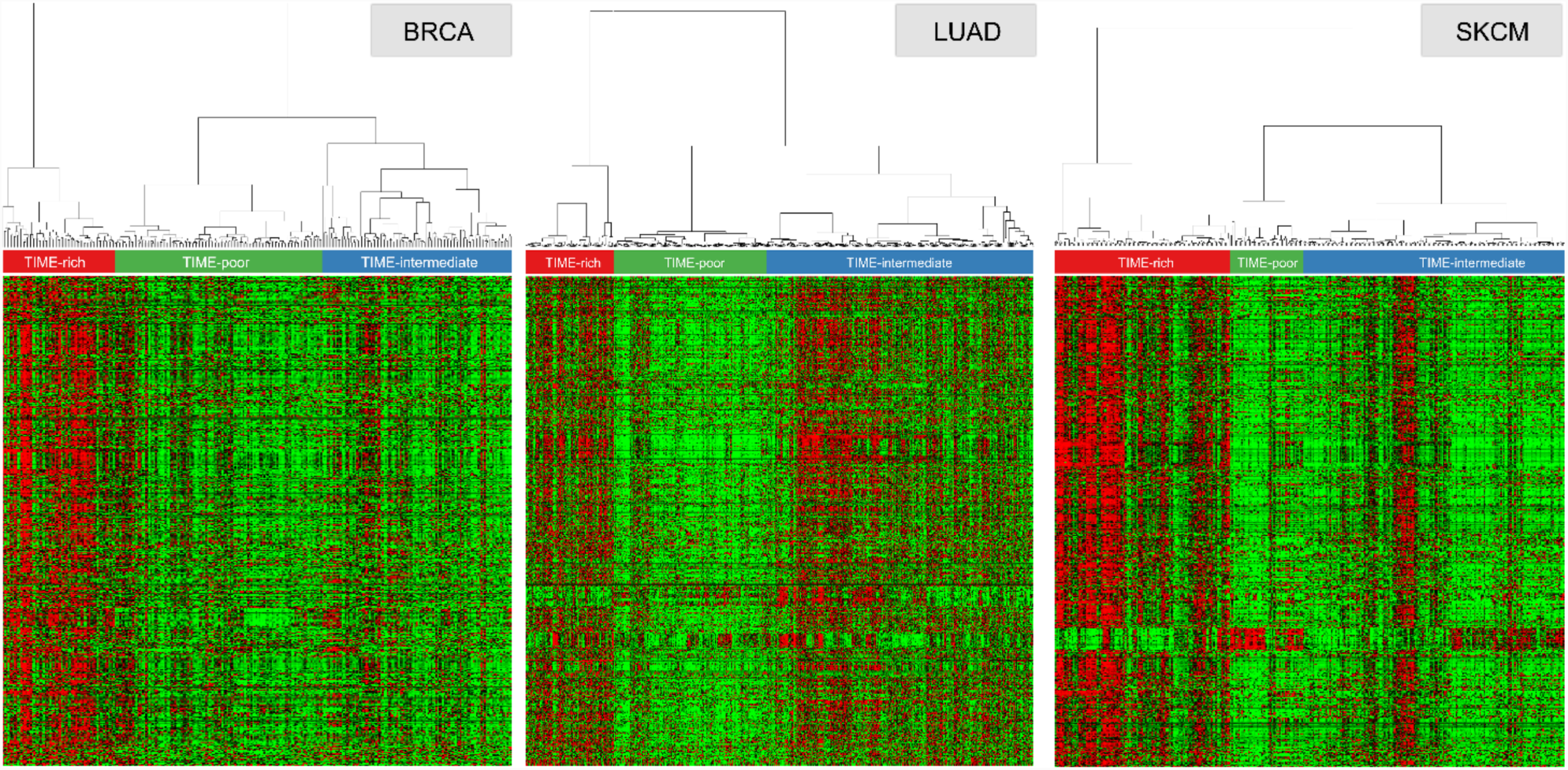
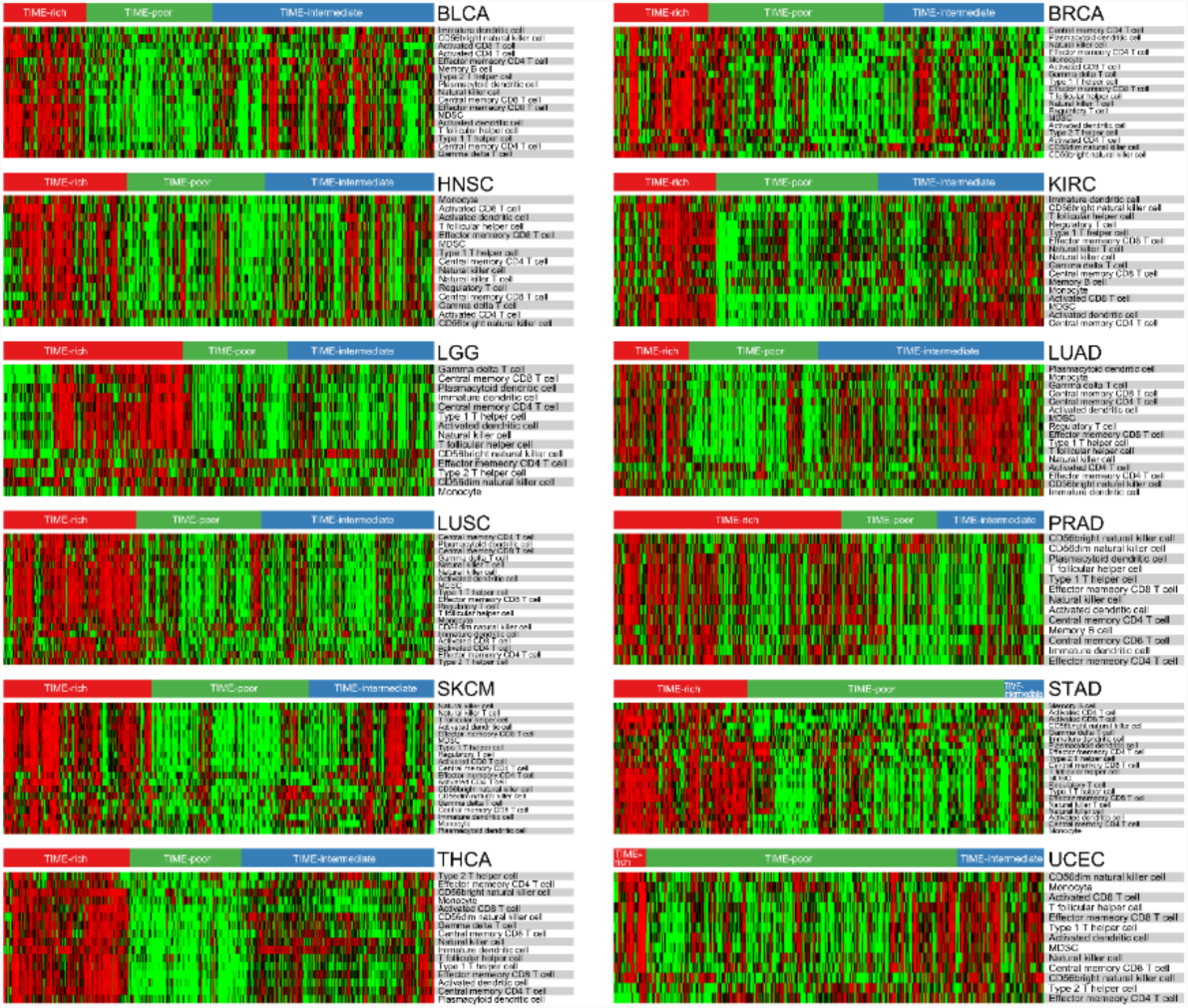
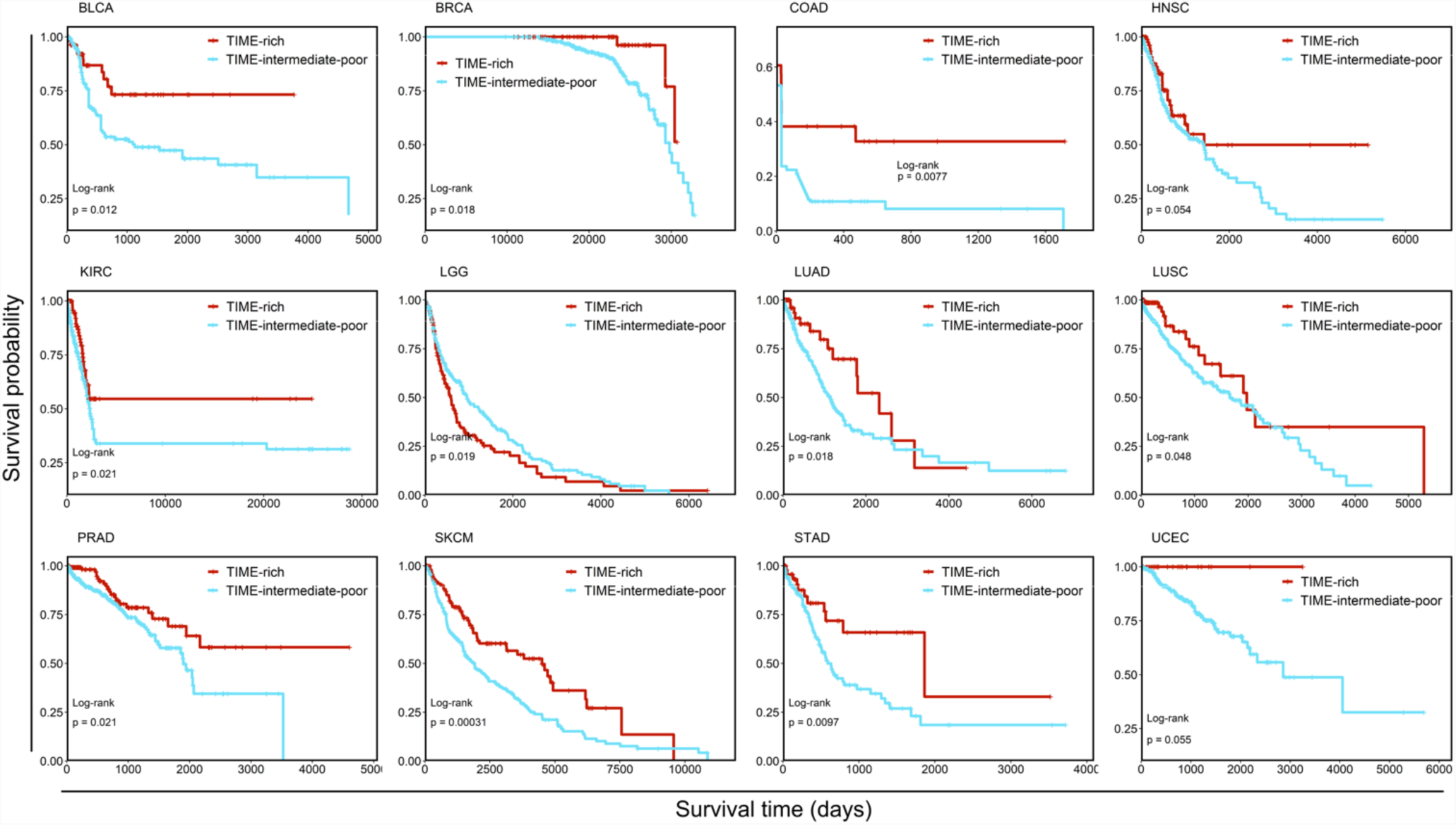
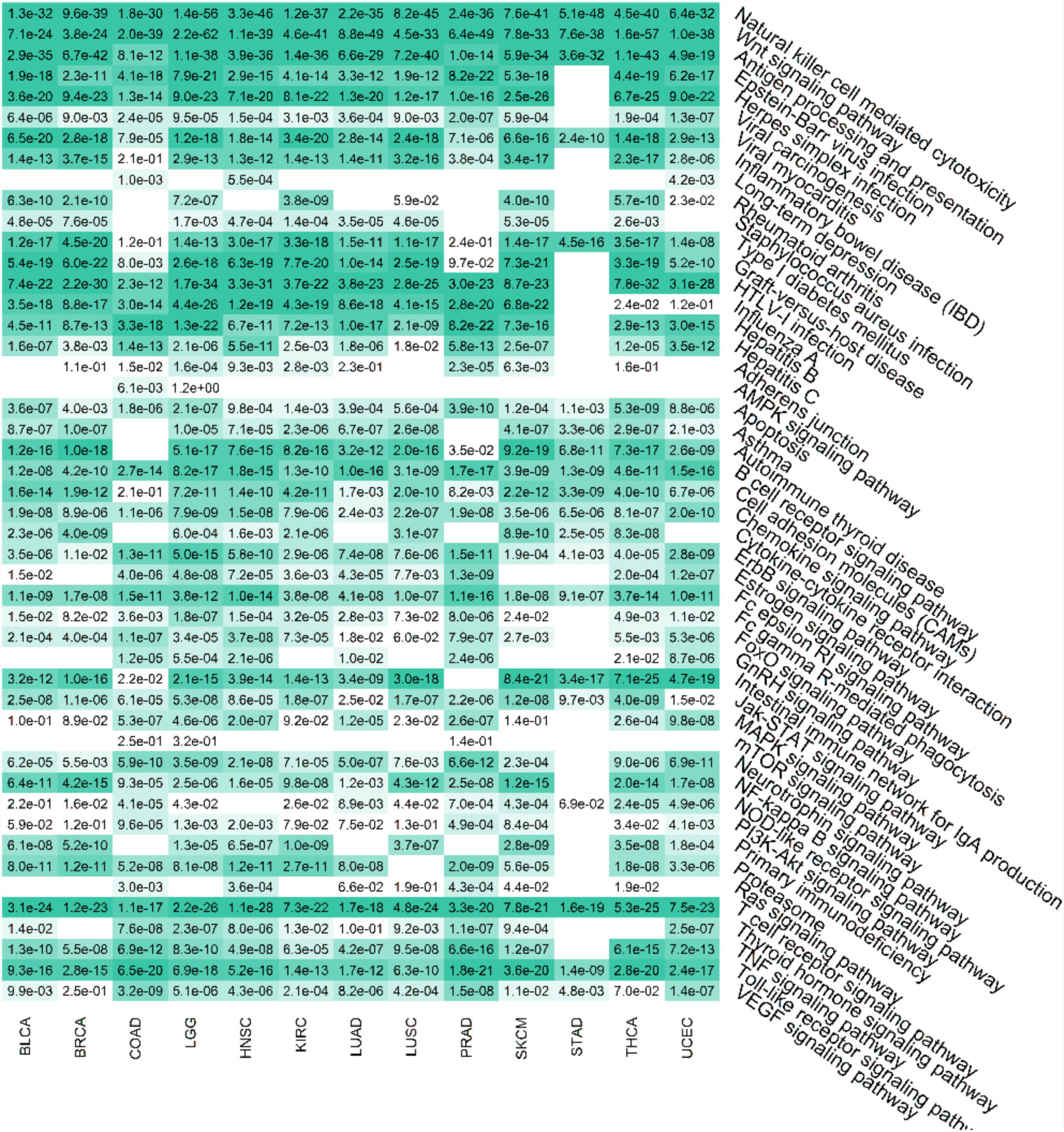
Clinical and cellular characteristics of the TIME subtypes. (a) Representative heatmaps derived from the gene expression of the immune-checkpoint therapy (ICT) essential genes, showing the universal TIME subtypes in breast invasive carcinoma (BRCA), lung adenocarcinoma (LUAD) and skin cutaneous Melanoma (SKCM). The heatmaps for other cancer types are shown in Supplementary Figure 1; (b) Abundance of infiltrated immune cells in TIME subtypes across 12 types of cancers; (c) Kaplan–Meier curves of the patient groups between the TIME-rich subtype and the combined TIME-intermediate and -poor subtypes, revealing that the survival time for the patients in the TIME-rich subtype is significantly longer than those in the TIME-intermediate and -poor subtypes; and (d) Pathway enriched analysis derived from the significantly differential genes of RNA-seq data between the TIME-rich subtype and the TIME-intermediate and -poor subtypes. The digital numbers represent FDR-corrected p values.

To discover the common features and critical differences which defined the 3 distinct TIME subtypes, we compared their gene expression profiles, TILs’ abundance in TIMEs and clinical outcomes. The abundance of TILs in TIMEs were significantly decreased from TIME-rich (‘immune-hot’ tumors), -intermediate (‘immune-cold’ tumors) to TIME-poor subtype (‘immune-desert’ tumors) (Fig 1b, Supplementary Table 1). It has been known that TILs are associated with tumor prognosis, thus we hypothesized that clinical outcomes could be different between TIME subtypes. In fact, both TIME-poor and -intermediate tumors had significantly poorer clinical outcomes than TIME-rich tumors (Fig 1c), however, patient survival differences were not significant between TIME-poor and -intermediate tumors. Pathway enrichment analysis of the significantly modulated genes between subtypes showed that the degree of the activated (i.e., represented by gene expression of each pathway or program) immune programs such as APP, NK cell-mediated cytotoxicity, T cell receptor singling pathways were significantly decreased from TIME-rich, -intermediate to -poor subtype (Fig 1d and Supplementary Fig 2). These results suggested that TIME-poor and -intermediate tumors had a lower degree of activated immune programs so that they could have a better chance to escape immunosurveillance. For example, APP is an immunological process that prepares and presents antigens to T cells. NK effector and T cell receptor signaling pathways function as tumor cell killing. Therefore, lower activity of APP, NK cells and T cell receptor singling allow tumor cells to have fewer immune constraints. These results were repeatedly observed across the 13 common cancer types, suggesting that the TIME subtypes are not only universal subtypes across cancer types but also share common cellular and clinical features.

In general, TIME-rich, -intermediate and -poor tumors represent 25.35%, 32.94%, and 41.70% of the tumors, respectively, depending on cancer types (Supplementary Table 2). These results indicated that TIME-rich patients represent only a small fraction of tumors, and more than 70% of the tumors are either TIME-poor or -intermediate tumors, most of which are known to non-respond to ICT. These results explain why only 10-40% of tumors (i.e., depending on the cancer type) respond to ICT.

### Inherited defects in NK cells shape TIME subtypes and clinical outcomes

The discovery of the universal TIME subtypes across the 13 common cancer types triggered us to hypothesize that the TIME subtypes could be modulated by common genetic regulators. We previously proposed that pre-existing inherited variants in germline genomes of cancer patients could play an important role in shaping somatic mutations, CNVs and oncogenic pathways in their paired tumor genomes (^15^). Furthermore, we have shown that inheritably functional variants of breast cancer patients significantly predicted tumor recurrence (^16, 17^), and the risk for breast, brain and other cancers (^18^). Moreover, inheritably functional variants of patients could be used to predict key somatic-mutated genes in their paired tumor genomes (Zaman et al., unpublished data). Finally, cancer patients’ germline genomes contain specific genomic patterns which are associated with cancer risk and clinical outcomes (Feng et al., unpublished data). These results raised a question whether cancer patients’ germline genomes modulated TIME subtypes.

To answer this question, we compared the functional germline variants (termed as inherited defects here, see Methods) of the patients between TIME subtypes and showed that genetic makeups of the patients were significantly different between them (Supplementary Fig 3): TIME-poor/-intermediate and TIME-rich patients had significantly differentially inherited genetic defects. Further pathway enrichment analysis showed that more than 30% of the defected or modulated pathways were associated with NK cell deficient phenotypes, NK cell-associated virus infections, and the NK cell-mediated cytotoxicity pathway (Fig 2a, Supplementary Fig 4). For example, among the significantly differential pathways and phenotypes, 7 of them were known NKD (NK cell deficiency) phenotypes (^19, 20, 21^) such as Epstein-Barr virus (EBV) infection, Herpes simplex infection, Leishmaniasis, Rheumatoid arthritis, Type I diabetes mellitus, long-term depression and viral myocarditis, while 7 of them were NK cell-related virus infections such as Graft-versus-host disease, HTLV-I infection, Hepatitis B and C, Influenza A, Asthma and IBD (Fig 1d, 2a, and 2b). These results strongly suggested that TIME-poor/-intermediate patients might have more inherited defects in their NK cells than TIME-rich patients.

**Figure 2.**
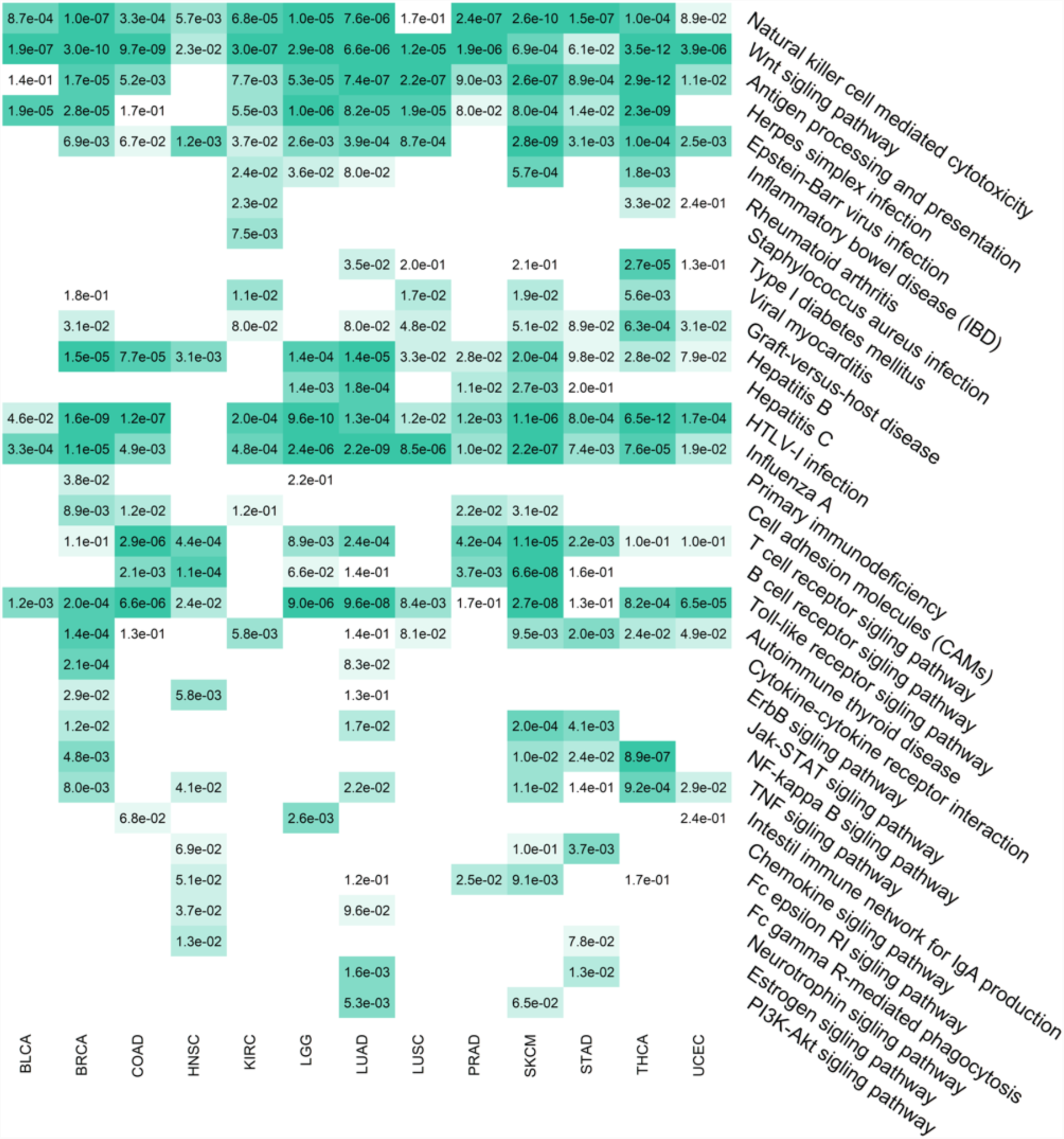
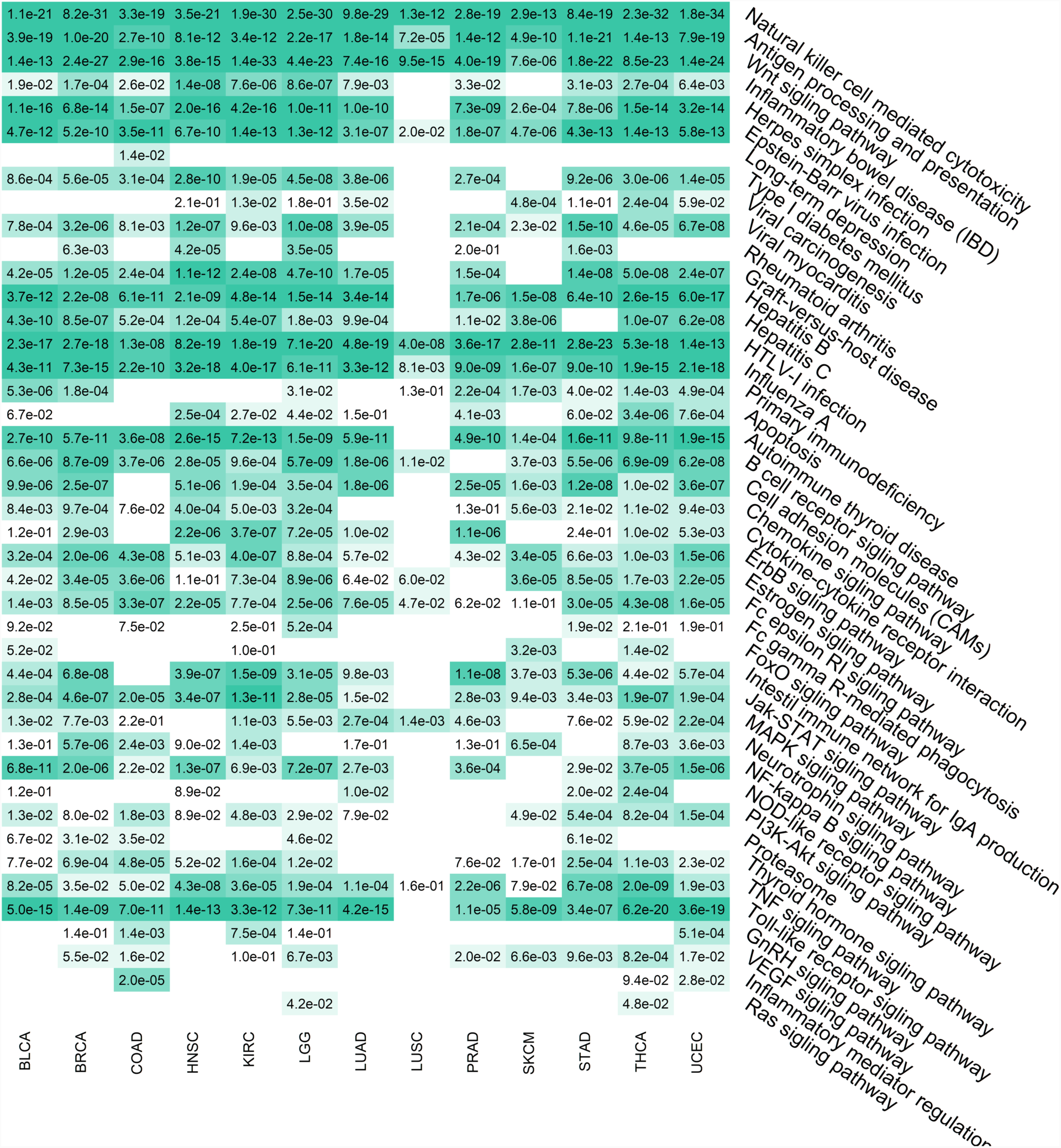
Heatmaps representing the enrichment pathways derived from functional germline genomic variants. (a) A heatmap shows the significantly enriched pathways derived from the significantly differential germline variants between TIME-rich and TIME-intermediate-poor subtypes in 13 cancer types. (b) A heatmap shows the significantly enriched pathways derived from the significantly differential germline variants between non-cancer individuals and TIME-intermediate-poor patients in 13 cancer types. The digital numbers represent FDR-corrected p values.

NK cells are the ‘first-line immune cells’ which enable to quickly detect and attack tumor cells and viruses. NK cells are able to control tumorigenesis by accurately regulating the distinct germline encoded inhibitory and activating NK cell surface receptors. In general, an excess of activating over inhibitory signals triggers the production and release of effector molecules, which can lead to the death of the infected or transformed cancer cell (^22^). Thus, to further explore the association between NK cell inherited defects with cancer progression and metastasis, we compiled a comprehensive set of NK cell genes including NK cell receptors and ligands, genes in NK cell signaling pathways (i.e., KEGG NK cell-mediated cytotoxicity pathway (^23, 24, 25^), and then conducted Fisher’s tests to identify genes which had significantly differential frequencies between TIME-rich group and TIME-poor/-intermediate group. For a given cancer type, we found 15-20 defected NK cell genes which had significantly higher frequencies in TIME-poor/-intermediate group than TIME-rich group (FDR-corrected p <0.25). The gene list for each cancer type was different, which could be associated with the diversity of tissue-resident NK cells, although many of the significantly defected NK cell genes of the 13 cancer types were shared. In total there were 103 such genes (i.e., termed as NK-genes here), 70 of which appeared in at least two cancer types. Among the 70 NK-genes, 37.2%, 25.8%, 24.3%, and 12.7% are known NKD genes (^19, 20, 21^), NK cell receptors and ligands, NK cell singling genes but also expressed in cell signaling in T or other immune cells (i.e., mainly innate immune cells, termed as ‘global immune cell genes’ here) and universal genes (i.e., expressed in many cell types), respectively (Supplementary Tables 3 and 4).

We further asked if inherited defected NK-genes were correlated with clinical outcomes. To answer this question, for each cancer type, we top-to-bottom ranked the patients based on the number of inheritably defected NK-genes. We defined top 40% and bottom 40% of the ranked patients as high- and low-NK cell defected groups, respectively, to conduct survival analysis. In 10 of the 12 cancer types (i.e., no survival data were available for THCA) except LGG and KIRC, the high-NK cell defected group had significantly shorter survival than the low-NK cell defected group (Fig 3a). Similar results were obtained when using top 30% and bottom 30% of the ranked patients as high-and low-NK cell defected groups, respectively. These results suggested that NK cells inherited defects could play an important role in tumor recurrence and overall survival in most of the cancer types. Given the fact that TILs’ abundance was different between TIMEs, we hypothesized that NK cell defected gene number could be negatively correlated with the TILs’ abundance in TIMEs. Indeed, it was true that for all the 13 cancer types (Fig 3b, Supplementary Fig 5), the abundance of immune cell troops such as NK, NKT, CD103+ DC (cDC), activated CD8+ T, CD4+ T, activated B, all kinds of T cells and 23 other types of immune cells were negatively correlated with the defected gene number in NK cells, suggesting that NK cells could be an important factor in recruiting other immune cells into TIMEs.

**Figure 3.**
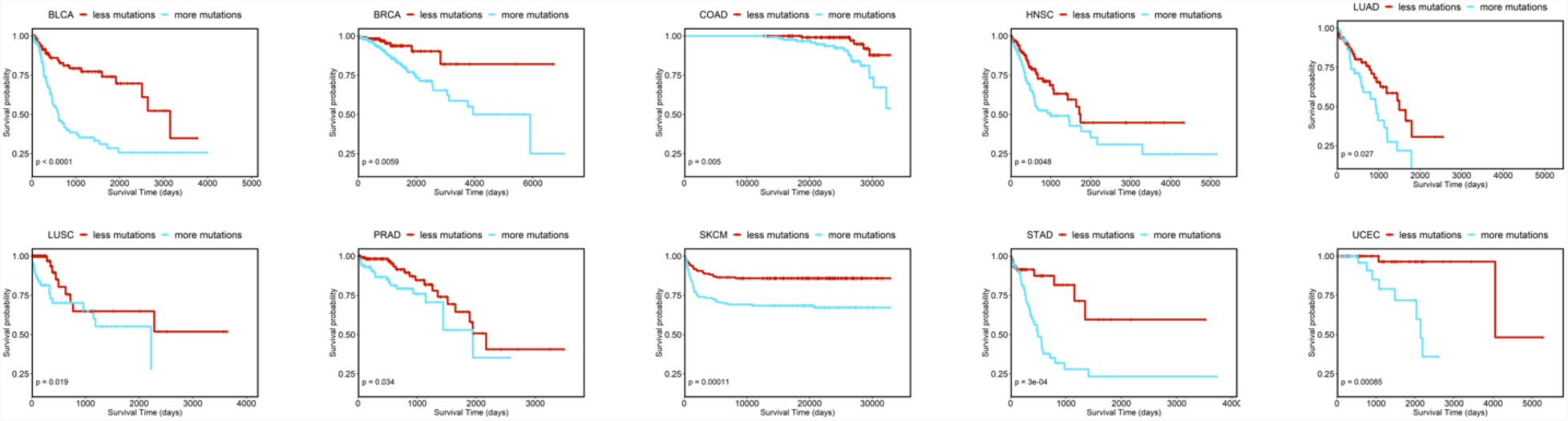
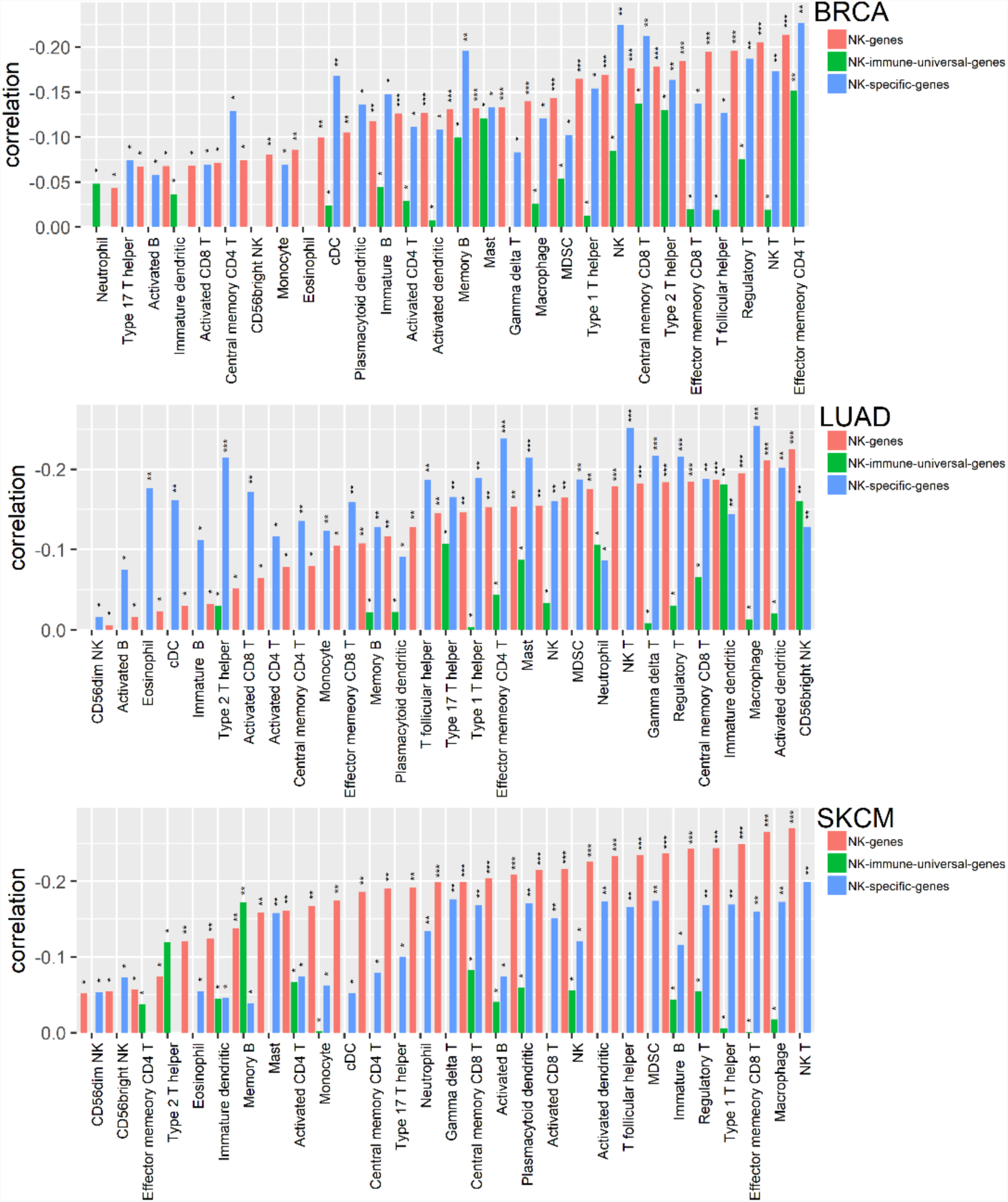
The associations between the inherited detected NK cells and clinical outcomes, and the abundance of infiltrated immune cells in TIMEs. (a) Kaplan–Meier curves of the patient groups of the high-and low-number of functionally inherited variants in the NK cells for disease-free survival. Patients were top-to-bottom ranked based on the number of functionally inherited variants in NK cells. Top 40% and bottom 40% of the ranked patients were defined as high-and a low number of the NK cell defected patient groups, respectively. (b) Negative correlations between the number of the NK cell inheritable defected and the abundance of the infiltrated immune cells in TIMEs. Breast invasive carcinoma (BRCA), lung adenocarcinoma (LUAD) and skin cutaneous Melanoma (SKCM). The survival differences for other types of cancer are shown in Supplementary Figure 6. *p-value >0.05; **0.05>p-value >0.001; and ***p-value<0.001. The NK-specific-genes included NKD genes and NK receptors/ligands of the 103 defected NK cell genes, while the NK-immune-universal-genes included the global immune cell genes and the universal genes of the 103 defected NK cell genes.

NKD genes and NK cell receptors/ligands are more preferentially involved in either NK cell-mediated cytotoxicity, NK cell development or proliferation (^11, 26^), while the global immune cell genes could influence the immune response of not only NK cells but also other immune cells. On the other hand, the universal genes play important roles in cell signaling of not only NK cells but also other cell types including tumor cells. Therefore, NKD genes and NK receptors/ligands are more NK cell-specific (i.e., representing 63% of the NK-genes, termed as NK-specific-genes here), while the global immune cell genes (i.e., representing 24.3% of the NK-genes) and the universal genes (i.e., representing 12.7% of the NK-genes) are expressed in multiple immune cells and non-immune cell types, respectively. Thus, we raised a question whether the correlations observed in Fig 3a, 3b and Supplementary Fig 5 were truly from NK cell defected genes. By extending the same analyses in Fig 3a and 3b using only the universal genes, there were not any correlation as shown in Fig 3a and 3b, suggesting that inherited defects in the universal genes alone are insufficient to modulate either TILs in TIMEs or clinical outcomes. Next, we re-did the same analyses by combining of the universal genes and the global immune cell genes (i.e., representing 37% of the NK-genes, termed as NK-immune-universal-genes here) and the NK-specific-genes, respectively. For the NK-immune-universal-genes, correlations, as shown in Fig 3a (i.e., survival), were not observed, on the other hand, for the NK-specific-genes, such correlations were reproduced across the 10 cancer types. These results suggested that the NK-specific-genes are sufficient to shape clinical outcomes, but the NK-immune-universal-genes were not.

Similarly, significantly negative correlations between TILs’ abundance in TIMEs and the number of the inheritably defected NK-specific-genes were reproduced in 10 cancer types including BLCA, BRCA, KIRC, LAUD, LUSC, THCA, UCEC, HNSC, LGG, and SKCM (Fig 3b, Supplementary Fig 5). In particular, the correlations derived from the NK-specific-genes were much stronger than those derived from the NK-genes in 7 cancer types (i.e., BLCA, BRCA, KIRC, LAUD, LUSC, THCA and UCEC, Fig 3b, Supplementary Fig 5). On the other hand, weaker negative correlations (i.e., correlation co-efficiencies were 2-10 times less than those derived from the NK-genes) and weaker statistical significances (i.e., p values in the range of 0.01-0.05) were observed for some immune cells in some cancer types when using the NK-immune-universal-genes (Fig 3b, Supplementary Fig 5). These results suggested that inherited defects of the NK-specific-genes are sufficient to impair communications between NK cells and other immune cells and then block TILs’ recruitment into TIMEs. Of note, in COAD, STAD, and PRAD, both NK-specific-genes and NK-immune-universal-genes did not reproduce the negative correlation (i.e., for TILs in TIMEs) derived from the NK-genes. PRAD has dense stroma (^27^) which could need the whole NK gene set (i.e., NK-genes) for TILs’ recruitment. NK cells surrounding both COAD and STAD are directly interacting with environmental factors such as microbiome, food, drinks, and others so that NK cells associated with both COAD (colon cancer) and STAD (gastric cancer) are much more complex. These data highlighted that inherited defected in NK cells are one of the key genetic factors in shaping TIME subtypes, TILs’ abundances and clinical outcomes.

Based on these insights, we hypothesized that NK cell was a critical factor for recruiting immune troops into TIMEs. This hypothesis is partially supported by two recent studies which had no idea about NK cell inherited defects (^27, 28^), but showing that depletion of NK cells resulted in failed recruitment of CD8+ T cells to the tumor microenvironment in melanoma mice. They further showed that NK cells recruited CD103+ DC, which in turn were required for the recruitment of effector T cells. Our results here suggested that NK cells could be able to recruit not only CD103+ DC (cDC) and CD8+ T cells but also 30 other immune cells. Because NK cells act as first responders to detect and kill cancer cells, it is reasonable to believe that NK cells could sequentially recruit many other immune cells including CD103+ DC and T cells into the tumor microenvironment. Thus, NK cells could recruit a whole immune troop but not only one or two immune cell types into TIMEs.

This insight provides a potential opportunity that adoptive cell transfer of NK cells from healthy donors could convert a TIME-poor tumor into a TIME-rich tumor. Adoptive NK cell was safe and does not cause a graft-versus-host disease attack on the recipients. Thus, we hypothesized that adoptive NK cell transfer of healthy donors (i.e., allogeneic) to a patient could have better clinical outcomes than the adoptive transfer of NK cells from the patients themselves (i.e., allogeneic) because cancer patients’ own NK cells are defected and don’t function very effectively. This hypothesis is supported by a recent clinical trial showing that allogenic adoptive NK cell transfer was much better than autogenic NK cell immunotherapy for breast cancer outcomes (^29^). TIME-rich tumors are more suitable to conduct immunotherapy than TIME-poor tumors, while TIME-rich tumors can gain significantly more survival benefit than TIME-poor tumors for neoadjuvant chemotherapy as well. For example, a meta-analysis of 13 studies showed that higher abundance of TILs in pre-treatment tumor biopsy was correlated with better pathological complete response rates to neoadjuvant chemotherapy(^30, 31, 32, 33, 34^). Therefore, it is a great interest to improve patient outcomes by applying adoptive transfer of healthy, functional NK cells to convert TIME-poor tumors into TIME-rich tumors which could be in turn treated with chemotherapy or immunotherapy. A recent clinical trial partially supported this hypothesis: adoptive transfer of NK cells in combination with chemotherapy in stage IV colon cancers significantly prevented recurrence and prolonged survival than chemotherapy alone. Most importantly, they found that poorly differentiated colon cancers were more susceptible to adoptive NK cell transfer than well-differentiated ones (^35^). Poorly and well-differentiated cancers are shorter and longer survival patients, respectively, which are corresponding to TIME-rich and -poor tumors, respectively. Therefore, these results agree with our above hypothesis. Another study also supported this hypothesis: the combination therapy of anti-PD-1 antibodies and activated autologous NK cells significantly delayed tumor progression in glioma-bearing animals as compared to the monotherapy regimens of anti-PD-1 or stimulated NK cells alone (P<0.001)(^36^).

### Tumorigenesis and metastasis are not random events and cancer patients are highly selected

As shown in Fig 2a and Supplementary Fig 4, except NK cell inherited defects, TIME-poor/-intermediate patients had significantly more defected APP and Wnt pathway genes than TIME-rich patients. APP is a biological process which presents antigens to T cells, functional defects in APP could lead tumor cells to escape T cell surveillance. However, careful analyses showed that APP pathway genetic defects were not significantly associated with clinical outcomes and TILs’ abundance. These results suggested that APP pathway defects alone were not sufficient to shape either clinical outcomes or TIME subtypes. In addition, it has been shown that activated Wnt signaling pathway in tumors excluded the recruitment of T cells into TIMEs (^37^). Furthermore, somatic mutations of the Wnt pathway in tumors could activate the pathway to prevent T cells to be recruited into TIMEs (^38^). Here we showed that in most of the cancer types the number of inherited defects in Wnt pathway in germline genomes had a positive correlation with the gene expression of the Wnt pathway (i.e., pathway activation) in their paired tumors, suggesting that a positive correlation between inherited defects in the Wnt pathway and activation of the pathway, which is similar to the somatic mutations in genes of the pathway.

The number of the inheritably defected genes in the Wnt pathway had weakly negative correlations (i.e., with marginally significant p values in the range of 0.02-0.05) with clinical outcomes in 8 cancer types (i.e., BLCA, BCRA, HNSC, LAUD, LUSC, PRAD, STAD and UCEC) and no such correlation with other cancer types (Supplementary Fig 6). Similarly, Wnt pathway inheritably defected gene number had negative correlations with the abundance of TILs in TIMEs (Supplementary Fig 7) in eight cancer types (i.e., HNSC, KIRC, LGG, LUAD, LUSC, SKCM, STAD and THCA), however, the correlations were weaker (i.e., in terms of correlation co-efficiencies and correlation significance represented by p values) than those derived from NK cells. These results suggested that inheritably defected in Wnt pathway could shape TILs’ recruitment in some cancer types, and the influences were much weaker than NK cells. Next, we asked if inheritably defects of the Wnt pathway and NK cells worked in a synergy manner. To answer this question, we conducted the correlation analysis using the genes combined from the NK cells and the Wnt pathway and showed that negative correlations between the inheritably defected gene number and the TILs’ abundance in TIMEs remained in eight cancers but much weaker than the correlations derived from either NK cells or Wnt pathway alone. These results suggested that both NK cells and Wnt pathway were parallel and complementary modulators to shape TILs recruitment, implying that to reverse immune exclusion in TIMEs, Wnt signaling inhibitors could be better than adoptive NK cell transfer in some tumors but not most of the tumors.

Together with the discovery that TIME-rich tumors have significantly longer survival than TIME-poor/-intermediate patients, we concluded that inherited defects in NK cells and Wnt pathways were associated with metastasis and clinical outcomes. These results highlighted that metastasis is affected by inherited defects and is not a random process. Similarly, significantly more inherited defects in NK cells, APP and Wnt pathways were observed between non-cancer individuals and cancer patients in the 13 common cancer types (Supplementary Fig 8). They were even more significant between TIME-poor/-intermediate patients and non-cancer individuals (Fig 2b and Supplementary Fig 9). These results suggested that individuals who bear inherited defects in NK cells, APP and Wnt pathways have a higher risk to get cancer. This hypothesis is indirectly supported by a study showing that individuals with lower NK cytotoxic activity in peripheral blood had a higher incidence of cancer (n=3,500 with 11-year follow-up)^39^). Similarly, previous studies showed that impaired NK cell activity was found in family members of patients with different types of cancer (^40, 41, 42^). Importantly, such inherited defects in NK cells, APP and Wnt pathways were repeatedly observed between non-cancer and cancer individuals (Fig 2b, Supplementary Fig 9), between TIME-rich and TIME-intermediate/-poor patients (Fig 2a, Supplementary Fig 4, Fig 1d), suggesting that these individuals were highly selected for tumorigenesis and metastasis. The immune programs (defects in NK cells, APP and Wnt pathways) are carried from germline genomes to their paired tumors. In fact, the number of inherited defects in the NK cells, APP and Wnt pathways was gradually increased from the non-cancer cohort, TIME-rich, -intermediate to -poor subtypes, suggesting that inherited defects in the NK cells, APP and Wnt pathways have a profound impact on cancer risk, TIME subtypes and clinical outcomes. These insights provided a potential opportunity to identify high-risk cancer subpopulation based on the inherited defects in NK cells, APP and Wnt pathways, further, adoptive NK cell transfer from healthy donors to high-risk individuals could postpone or prevent cancer development. This hypothesis is partially supported by a study showing that NK cell depletion in melanoma mice resulted in substantial metastasis, but the adoptive transfer of NK cells protects NK cell-deficient mice from tumor establishment (^43^).

In addition, as shown in Fig 2b, Type I diabetes and long-term depression phenotypes were linked to some cancers, suggesting that they were genetic diseases as well. Diabetes and obesity have been shown to be a risk factor for some cancers, for example, a meta-analysis of 121 cohorts including 20 million individuals and one million events confirmed that diabetes is a risk factor for all-site cancer (^44^), while obesity can increase cancer incidences of 13 cancer types (^45^). A 24-year follow-up study showed that depression increases the risk of cancer (^46^), Moreover, a meta-analysis of 16 studies (n=163,000) showed that cancer patients with anxiety and depression had a greater risk of dying from all types of cancer (^47^). Impairing of the NK cell function is one of the common factors behind these links. For example, obesity has been known to impair NK cell function and then lead to an increased risk for severe infections and several cancer types (^48, 49^), while chronic family stress is consistently associated with decreases in NK cell cytotoxicity (^50^). These results indicated that NK function impaired by either genetic defects or regulatory factors such as depression, obesity and diabetes could increase cancer incidence.

### TIME subtypes could guide in immunotherapies

As shown above, both NK cell-mediated cytotoxicity and Wnt signaling pathways became the hallmarks which enabled to significantly distinguish TIME-rich and TIME-intermediate/-poor subtypes. From the ICT clinical trials (^51,52^), we obtained 49 melanoma (SKCM, 10 responders) and 47 gastric (STAD, 12 responders) samples which had tumor RNA-Seq data before administrating of ICT. We used the significantly modulated genes of both pathways between TIME subtypes to assign the ICT-clinical trial samples into either TIME-rich or TIME-intermediate/-poor group based on the k-nearest neighbor algorithm (KNN, see Methods). By doing so, 70% and 30% of the ICT-responding melanoma were assigned into TIME-rich and TIME-intermediate/-poor group, respectively. In contrast, 56% and 44% of the ICT non-responding melanoma were assigned into TIME-rich and TIME-intermediate/-poor group, respectively. Similarly, 58% and 42% of the ICT-responding gastric tumors were assigned into TIME-rich and TIME-intermediate/-poor group, respectively. In contrast, 31% and 69% of the ICT non-responding gastric tumors were assigned into TIME-rich and TIME-intermediate/-poor group, respectively. Not surprisingly, these results indicated that TIME-rich patients were enriched with ICT responders. Pathway enrichment analysis of tumor gene expression profiles showed that TIME-rich non-responders had significantly higher gene expression in Wnt pathway (FDR-corrected p=5.3×10^-12^ and 4.0×10^-5^ for melanoma and gastric tumors, respectively) than TIME-rich non-responders, suggesting that using of Wnt inhibitors could improve ICT treatment for TIME-rich non-responders. Similarly, genes of the NK cell-mediated cytotoxicity pathway (FDR-corrected p=2.5×10^-3^ and 2.7×10^-3^ for melanoma and gastric tumors, respectively) and APP (FDR-corrected p=7.1×10^-4^ for melanoma) were significantly lower expressed in TIME-intermediate/-poor non-responders than TIME-intermediate/-poor responders, suggesting that adoptive NK cell transfer in combination with CAR-T or ICT could improve the existing immunotherapies for TIME-intermediate/-poor non-responders. However, it should be cautious about these conclusions due to the small number of clinical trial samples (n=96) here, more samples are needed to further validate these discoveries in the future.

## Discussion

### A unified view of the tumor immune microenvironment

Tumor molecular subtypes could inform not only prognosis but also treatment. Here we showed that TIMEs can be classified into three universal subtypes across 13 common cancer types. Different from previous observations of the complex tumor molecular subtypes, each of which often has its own unique features, the universal TIME subtypes were commonly shared by multiple cancer types, providing a framework to get better insight into the unifying features of each TIME subtype and the differences between the distinct TIME subtypes, and then understand why some tumors respond to immunotherapy and some don’t. Furthermore, this framework allows exploring genetic factors to regulate TIMEs and could help in identifying new druggable targets. Regardless of cancer types, TIME subtypes have unifying features: (1) TIME-rich patients have significantly longer survival than other TIME subtypes, because (2) the abundance of the TILs in TIME-rich, -intermediate and -poor tumors is gradually decreased. These features recaptured the known immune-hot, -cold and -desert tumors described previously based on immunohistochemistry (^53, 54^). It has been well-known that a lower abundance of TILs in TIMEs is associated with more cancer recurrence and shorter survival (^55, 56^). (3) signaling pathways associated with key immune programs such as APP, NK, and T cell signaling were more highly activated in TIME-rich tumors than TIME-intermediate/-poor tumors. The degree of the activated immune programs (i.e., represented by the expression levels of the pathway genes) was gradually decreased from TIME-rich, -intermediate to -poor tumors. Lower level expression of the genes in these immune-programs and pathways will allow tumors to escape from immune attack. (4) finally, ICT-responders were more enriched in TIME-rich tumors.

### Cancer is an NK cell deficient disease

The hypothesis of cancer immunosurveillance suggests that tumor cell transformation occurs frequently, but is under constant control by the immune system. The immune system is able to identify transformed cells that have escaped cell-intrinsic tumor-suppressor mechanisms and eliminate them before they can establish malignancy (^57^). Thus, if the individuals are genetically immunodeficient, they could have a markedly increased incidence of cancer. In general, NK cells have a large repertoire of germline-encoded inhibitory and activating receptors to sense ‘danger’ in the cell surfaces. The germline-encoded receptors which recognize ligands associated with viral infection or cancer cell transformation (^10^). Genetic defects in NK-genes including NK cell receptors and NKDs could impair NK cell functions. By conducting an unbiased scanning of germline genomes of cancer patients, we provided a clearer view of the spectrum of malignancies associated with impaired NK cells and showed that inherited defects affecting NK cell functions were sufficient to accomplish cancer immunosurveillance of the immune system.

Among highly related pathways (i.e., NK cells, APP and Wnt pathways) examined, only NK-genes’ defects were correlated with both survival and the abundance of TILs in TIMEs. Most importantly, in 10 out of the 12 common cancers (i.e., no survival data were available for THCA), defects of the NK-specific-genes (i.e., NKDs + NK cell receptors) alone were sufficient to establish these correlations (i.e., survival and TILs’ abundance). Traditionally, the hypothesis of cancer immunosurveillance mainly focuses on T cell or other immune cells for their cell-killing function. Here we demonstrated that to implement cancer immunosurveillance, NK cells could play a critical role in communicating with many immune cells to recruit the whole immune troops into the cancer-transformed cell or TIMEs. Specifically, our results suggested that NK cells could have two functions for controlling tumor progression and metastasis: (1) cancer cell killing (2) recruiting immune troops into TIMEs. Here we proposed a working model for NK cell-based immunosurveillance: NK cells are the first to arrive in the tumor microenvironment and recruit CD103+ DC through the secretion of chemokines. TIL-DCs then recruit effector T cells. Activated NK cells produce numerous cytokines, communicate with other 30 immune cells, and then recruiting the immune troops into the cancer cell or TIMEs. Anti-tumor and anti-metastasis activity (i.e., survival differences between TIME subtypes) is thus the result of the collaboration of distinct innate and adaptive immune cell types. Thus, inheritably defected NK-genes could impair NK cell function and then block to recruit the immune troops to the cancer cell or into TIMEs to conduct the anti-tumor activity. Thus, we proposed that cancer is largely a disease of NK cell deficiencies. This working model unraveled the cellular and molecular determinants (i.e., NK cells) of multiple immune cell recruitment to and retention in solid tumors’ TIMEs. Thus, allogeneic adoptive NK cell therapy (i.e., collecting healthy NK cells which have no inherited defects from donors and infusing them into the patient) could convert immunotherapy-resistant, TIME-poor tumors into ICT/CAR-T-sensitive, TIME-rich tumors. This NK cell therapy could be a universal approach which is independent of cancer types. Given the fact that more than 70% of the cancer patients are immune-cold and -desert subtypes, successfully validating of this hypothesis will be a crucial step toward improving the existing immunotherapies.

In addition, our results suggested that inherited defects in NK cells, APP and Wnt pathways were positively selected in tumorigenesis and metastasis. Therefore, cancer patients are highly selected, and thus, it provides an explanation for the fact that only 12-20% of the heavy smokers develop lung cancer in their lifetime, although 85% of the lung cancer patients are heavy smokers (^58, 59^). Cancer causally environmental factors play an important role in tumorigenesis, however, without a susceptible germline genome (i.e., genetic defects in NK cells, APP and Wnt pathways), they still cannot induce cancer. We proposed that germline genome is the most important factor in determining if a person will get cancer in one’s lifetime, and cancer is the result of the interactions between high-risk germline genomes and risk-factors (i.e., environmental factors and lifestyles).

Thus, the discoveries in this study could open a new window to explore NK cell biology and lead to novel thinking about identifying high-risk individuals for early cancer detection, precision cancer prevention, and immunotherapy. Strategies to harness and augment NK cells for cancer therapy are relatively new and rapidly developing field and have not been used for cancer prevention yet. The concept that cancer is an NK cell deficient disease could lead this field to new directions: (1) identifying of high-risk population based on NK cell, APP and Wnt pathways’ inherited defects, so that early cancer detection or precision prevention (i.e., by avoiding exposure to smoking, UV lights and other risk factors) could be implemented. (2) Preventing or postponing of cancer development for the high-risk population by adoptive NK cell transfer. (3) Converting TIME-poor/-intermediate tumors into TIME-rich tumors by manipulating NK cells, adoptive NK cell transfer or using Wnt signaling inhibitors so that ICT or CAR-T therapy could be applicable. Finally, many open questions still remained, for example, defected genes in NK cells were largely shared by different cancer types, however, each cancer type has some unique NK cell gene defects. Given the fact that tissue/organ-resident NK cells are different and diverse (^60^), additional studies will be needed to elucidate if defected genes are dominantly expressed in tissue/organ-resident NK cells. If so, adoptive transfer of tissue-resident NK cells could be considered to be more efficient in cancer prevention and TIL’s recruitments. NK cells share lots of expression programs with NKT cells, which is a subset of innate-like T cells. Both NK cells and NKT cells are cytotoxic cells, which trigger innate immune responses, provide the first level of defense against infected cells and tumor cells, produce cytokines and trigger immune responses without a prior sensitization by the immune system. Along with this long, we suspected that the inherited defects in NKT cells could also play similar roles which were discussed in this study, although the cell number of the NK cells is nearly 200 times of the NKT cells.

## Methods

### Datasets

Based on the availability of the whole exome sequencing (WES) of germline gnomes, their paired tumor genomes, and paired RNA-seq data, 14 common cancer types were selected from TCGA (n=200 at least for each cancer type). To remove the effects of virus-infection for the NK cell study, we removed the virus-infected tumors. Only 15% of the liver tumors (i.e., this number is too small to conduct analysis) without virus infection, thus, we excluded this cancer type for further analysis. Thus, 13 common cancers used in this study were BLCA, BRCA, COAD, LGG, HNSC, KIRC, LUAD, LUSC, PRAD, SKCM, STAD, THCA, and UCEC. Because the primary melanoma samples were not many, only metastatic SKCM samples were included for analysis. Distinct clinical subtypes have been reported in breast cancer, thus we took only the ER+ breast tumors in this study because the sample numbers of HER2+ and triple negatives are small in TCGA. The WES files derived from buffy coats (normal samples) of these cancer patients (n=5,883) were downloaded from TCGA, and the normalized RNA-sequencing data with Root Mean Square Error (RMSE) across 13 types of cancers (n=5,373) were downloaded from FireBrowse. The WES files of 4,500 non-cancer individuals (phs000473.v2.p2, phs000806.v1.p1, phs001194.v2.p2) were downloaded from The database of Genotypes and Phenotypes (dbGaP). RNA-sequencing data of 49 ICT-clinical trial melanoma sample and 47 ICT-clinical trial gastric samples were collected from GSE91061 and PRJEB25780, respectively, and were processed based on the mRNA analysis pipeline in TCGA.

### Variant calling and germline variant determination

For TCGA WES files, variant calling was performed using Varscan (version 2.3.9). The thresholds for germline variant calls required variant allele fraction (VAF) between 45% and 55%, and >90%. Functional variants were examined and annotated using the Combined Annotation Dependent Depletion (CADD) using the default parameters. To keep the consistency with the TCGA pipeline for processing WES data of the non-cancer individuals, BWA (version 0.7.15) was used to align with default parameters, piping into Samtools (version 0.1.8) to sort. Additional adding read groups and duplicate removal were processed with Picard-tools (version 2.6.0). The resulted BAM files were processed with GATK (version 4.0.11.0) for realignment and base recalibration. Varscan (version 2.3.9) and CADD were used to call and annotate functional germline variants.

### Immune gene set and clustering analysis

Immune-related genes (n=1,384) including MHC system-related genes [^61^], immunophenoscore-related genes [^14^], ICT essential genes for immunotherapy [^12^] and cytotoxic T cell-resistant genes [^62^] were collected and identified as critical immune-related genes (the gene pool *G)*. RNA-sequencing data of melanoma patients in TCGA were used to screen the key genes.

Step#1 Initialize the candidate set of key genes, that is, *G*_*candidate*_ = *ϕ*

Step#2 Randomly select 30% genes fro *G* m the gene set *G*_*random*_.

Step#3 Replace the features of elements in the patient set *P* with *G*_*random*_ to form the sample set *S*_*random*_.

Step#4 Group the samples *S*_*random*_ by using the hierarchical clustering method. For each of the clustering, clValid (^63^) was used to evaluate the clustering stability and the most stable clustering number was recorded.

Step#5 Repeat Steps 2-4 100,000 times. Rank the most stable clustering numbers and select the most suitable clustering number 3.

Step#6 Extract the genes when clustering number is 3, and rank the genes. Select the most informative genes and record them as the final set of key-gene candidates (1,294 genes).

We used the 1,294 genes to conduct unsupervised clustering analyses of the RNA-seq data for each cancer type to define TIME subtypes.

### TIL abundance calculation and pathway enrichment analysis

A deconvolution approach (^14^) was used to extract the abundance of each immune cell based on their gene markers from a tumor RNA-Seq data. Pathway enrichment analysis was conducted using DAVID (https://david.ncifcrf.gov).

### Assigning ICT trial samples into TIME subtypes

To assign an ICT-clinical trial sample into a TIME subtype, t-test statistics were first conducted in TIME-rich vs TIME-intermediate and TIME-intermediate vs TIME-poor tumors’ RNA-seq data for the genes derived from the NK cell-mediated cytotoxicity pathway and the Wnt signaling pathway. The significantly differential genes (p<0.05) were used for conduct K-nearest neighbors (KNN, k=5) to assign the sample to a TIME subtype. Spearman’s rank correlation was conducted between the sample and the samples in each TIME subtype based on the differential genes. TCGA-SKCM and TCGA-STAD samples were used for assigning the ICT-clinical trial samples of SKCM and STAD, respectively.

## References

1. Fallahpour S, Navaneelan T, De P, Borgo A. Breast cancer survival by molecular subtype: a population-based analysis of cancer registry data. CMAJ open 2017, 5(3): E734–E739.

2. Gao JJ, Swain SM. Luminal A Breast Cancer and Molecular Assays: A Review. The oncologist 2018, 23(5): 556–565.

3. Zaman N, Li L, Jaramillo ML, Sun Z, Tibiche C, Banville M, et al. Signaling network assessment of mutations and copy number variations predict breast cancer subtype-specific drug targets. Cell reports 2013, 5(1): 216–223.

4. Haque R, Ahmed SA, Inzhakova G, Shi J, Avila C, Polikoff J, et al. Impact of breast cancer subtypes and treatment on survival: an analysis spanning two decades. Cancer epidemiology, biomarkers & prevention : a publication of the American Association for Cancer Research, cosponsored by the American Society of Preventive Oncology 2012, 21(10): 1848–1855.

5. Sharma P, Allison JP. The future of immune checkpoint therapy. Science 2015, 348(6230): 56–61.

6. Postow MA, Callahan MK, Wolchok JD. Immune Checkpoint Blockade in Cancer Therapy. J Clin Oncol 2015, 33(17): 1974–U1161.

7. Peng D, Kryczek I, Nagarsheth N, Zhao L, Wei S, Wang W, et al. Epigenetic silencing of TH1-type chemokines shapes tumour immunity and immunotherapy. Nature 2015, 527(7577): 249–253.

8. Ribas A, Dummer R, Puzanov I, VanderWalde A, Andtbacka RHI, Michielin O, et al. Oncolytic Virotherapy Promotes Intratumoral T Cell Infiltration and Improves Anti-PD-1 Immunotherapy (vol 170, 1109.e1, 2017). Cell 2018, 174(4): 1031–1032.

9. Pahl J, Cerwenka A. Tricking the balance: NK cells in anti-cancer immunity. Immunobiology 2017, 222(1): 11–20.

10. Malmberg KJ, Carlsten M, Bjorklund A, Sohlberg E, Bryceson YT, Ljunggren HG. Natural killer cell-mediated immunosurveillance of human cancer. Seminars in immunology 2017, 31: 20–29.

11. Bryceson YT, Chiang SC, Darmanin S, Fauriat C, Schlums H, Theorell J, et al. Molecular mechanisms of natural killer cell activation. Journal of innate immunity 2011, 3(3): 216–226.

12. Patel SJ, Sanjana NE, Kishton RJ, Eidizadeh A, Vodnala SK, Cam M, et al. Identification of essential genes for cancer immunotherapy. Nature 2017, 548(7669): 537-+.

13. Rock KL, Reits E, Neefjes J. Present Yourself! By MHC Class I and MHC Class II Molecules. Trends Immunol 2016, 37(11): 724–737.

14. Charoentong P, Finotello F, Angelova M, Mayer C, Efremova M, Rieder D, et al. Pan-cancer Immunogenomic Analyses Reveal Genotype-Immunophenotype Relationships and Predictors of Response to Checkpoint Blockade. Cell reports 2017, 18(1): 248–262.

15. Wang E, Zaman N, McGee S, Milanese JS, Masoudi-Nejad A, O’Connor-McCourt M. Predictive genomics: a cancer hallmark network framework for predicting tumor clinical phenotypes using genome sequencing data. Seminars in cancer biology 2015, 30: 4–12.

16. Jean-Sebastien Milanese CT, Naif Zaman, Jinfeng Zou, Pengyong Han, Zhi Gang Meng, Andre Nantel, Arnaud Droit, Edwin Wang. Germline genomic landscapes of breast cancer patients significantly predict clinical outcomes. bioRxiv 2018.

17. Jean-Sebastien Milanese CT, Naif Zaman, Jinfeng Zou, Pengyong Han, Zhiganag Meng, Andre Nantel, Arnaud Droit, Edwin Wang. eTumorMetastasis, a network-based algorithm predicts clinical outcomes using whole-exome sequencing data of cancer patients. bioRxiv.

18. Jinfeng Zou EW. eTumorRisk, an algorithm predicts cancer risk based on co-mutated gene networks in an individual’s germline genome. 2018.

19. Orange JS. Natural killer cell deficiency. The Journal of allergy and clinical immunology 2013, 132(3): 515–525.

20. Mace EM, Orange JS. Genetic Causes of Human NK Cell Deficiency and Their Effect on NK Cell Subsets. Frontiers in immunology 2016, 7: 545.

21. Mandal A, Viswanathan C. Natural killer cells: In health and disease. Hematology/oncology and stem cell therapy 2015, 8(2): 47–55.

22. Long EO, Kim HS, Liu D, Peterson ME, Rajagopalan S. Controlling natural killer cell responses: integration of signals for activation and inhibition. Annual review of immunology 2013, 31: 227–258.

23. Watzl C, Long EO. Signal transduction during activation and inhibition of natural killer cells. Current protocols in immunology 2010, Chapter 11: Unit 11 19B.

24. Moretta A, Bottino C, Vitale M, Pende D, Cantoni C, Mingari MC, et al. Activating receptors and coreceptors involved in human natural killer cell-mediated cytolysis. Annual review of immunology 2001, 19: 197–223.

25. Biassoni R. Human natural killer receptors, co-receptors, and their ligands. Current protocols in immunology 2009, Chapter 14: Unit 14 10.

26. Abel AM, Yang C, Thakar MS, Malarkannan S. Natural Killer Cells: Development, Maturation, and Clinical Utilization. Frontiers in immunology 2018, 9: 1869.

27. Neesse A, Bauer CA, Ohlund D, Lauth M, Buchholz M, Michl P, et al. Stromal biology and therapy in pancreatic cancer: ready for clinical translation? Gut 2018.

28. Bottcher JP, Bonavita E, Chakravarty P, Blees H, Cabeza-Cabrerizo M, Sammicheli S, et al. NK Cells Stimulate Recruitment of cDC1 into the Tumor Microenvironment Promoting Cancer Immune Control. Cell 2018, 172(5): 1022–1037 e1014.

29. Liang SZ, Xu KC, Niu LZ, Wang XH, Liang YQ, Zhang MJ, et al. Comparison of autogeneic and allogeneic natural killer cells immunotherapy on the clinical outcome of recurrent breast cancer. Oncotargets Ther 2017, 10: 4273–4281.

30. Mao Y, Qu Q, Zhang Y, Liu J, Chen X, Shen K. The value of tumor infiltrating lymphocytes (TILs) for predicting response to neoadjuvant chemotherapy in breast cancer: a systematic review and meta-analysis. PloS one 2014, 9(12): e115103.

31. Denkert C, Loibl S, Noske A, Roller M, Muller BM, Komor M, et al. Tumor-associated lymphocytes as an independent predictor of response to neoadjuvant chemotherapy in breast cancer. J Clin Oncol 2010, 28(1): 105–113.

32. Oda N, Shimazu K, Naoi Y, Morimoto K, Shimomura A, Shimoda M, et al. Intratumoral regulatory T cells as an independent predictive factor for pathological complete response to neoadjuvant paclitaxel followed by 5-FU/epirubicin/cyclophosphamide in breast cancer patients. Breast cancer research and treatment 2012, 136(1): 107–116.

33. Miyashita M, Sasano H, Tamaki K, Chan M, Hirakawa H, Suzuki A, et al. Tumor-infiltrating CD8+ and FOXP3+ lymphocytes in triple-negative breast cancer: its correlation with pathological complete response to neoadjuvant chemotherapy. Breast cancer research and treatment 2014, 148(3): 525–534.

34. Azimi F, Scolyer RA, Rumcheva P, Moncrieff M, Murali R, McCarthy SW, et al. Tumor-infiltrating lymphocyte grade is an independent predictor of sentinel lymph node status and survival in patients with cutaneous melanoma. J Clin Oncol 2012, 30(21): 2678–2683.

35. Li LY, Cui JW, Wang C, Wang YZ, Niu C, Yao C, et al. Adoptive transfer of NK cells in combination with chemotherapy to improve outcomes of patients with locally advanced colon carcinoma. J Clin Oncol 2017, 35.

36. Multhoff MSEPBNLYYMIGBMG. P04.02 Adoptive transfer of lymphokine activated NK cells in combination with anti-PD-1 immune checkpoint inhibitor in therapy of glioblastoma. Neuro-Oncology, 20(Suppl_3): iii278.

37. Spranger S, Bao R, Gajewski TF. Melanoma-intrinsic beta-catenin signalling prevents anti-tumour immunity. Nature 2015, 523(7559): 231–235.

38. Grasso CS, Giannakis M, Wells DK, Hamada T, Mu XJ, Quist M, et al. Genetic Mechanisms of Immune Evasion in Colorectal Cancer. Cancer discovery 2018, 8(6): 730–749.

39. Imai K, Matsuyama S, Miyake S, Suga K, Nakachi K. Natural cytotoxic activity of peripheral-blood lymphocytes and cancer incidence: an 11-year follow-up study of a general population. Lancet 2000, 356(9244): 1795–1799.

40. Montelli TC, Peracoli MT, Gabarra RC, Soares AM, Kurokawa CS. Familial cancer: depressed NK-cell cytotoxicity in healthy and cancer affected members. Arquivos de neuro-psiquiatria 2001, 59(1): 6–10.

41. Strayer DR, Carter WA, Brodsky I. Familial occurrence of breast cancer is associated with reduced natural killer cytotoxicity. Breast cancer research and treatment 1986, 7(3): 187–192.

42. Bovbjerg DH, Valdimarsdottir H. Familial cancer, emotional distress, and low natural cytotoxic activity in healthy women. Annals of oncology : official journal of the European Society for Medical Oncology 1993, 4(9): 745–752.

43. Ballas ZK, Buchta CM, Rosean TR, Heusel JW, Shey MR. Role of NK cell subsets in organ-specific murine melanoma metastasis. PloS one 2013, 8(6): e65599.

44. Ohkuma T, Peters SAE, Woodward M. Sex differences in the association between diabetes and cancer: a systematic review and meta-analysis of 121 cohorts including 20 million individuals and one million events. Diabetologia 2018, 61(10): 2140–2154.

45. Steele CB, Thomas CC, Henley SJ, Massetti GM, Galuska DA, Agurs-Collins T, et al. Vital Signs: Trends in Incidence of Cancers Associated with Overweight and Obesity - United States, 2005-2014. MMWR Morbidity and mortality weekly report 2017, 66(39): 1052–1058.

46. Gross AL, Gallo JJ, Eaton WW. Depression and cancer risk: 24 years of follow-up of the Baltimore Epidemiologic Catchment Area sample. Cancer causes & control : CCC 2010, 21(2): 191–199.

47. Batty GD, Russ TC, Stamatakis E, Kivimaki M. Psychological distress in relation to site specific cancer mortality: pooling of unpublished data from 16 prospective cohort studies. Bmj 2017, 356: j108.

48. Bahr I, Jahn J, Zipprich A, Pahlow I, Spielmann J, Kielstein H. Impaired natural killer cell subset phenotypes in human obesity. Immunologic research 2018, 66(2): 234–244.

49. O’Shea D, Cawood TJ, O’Farrelly C, Lynch L. Natural killer cells in obesity: impaired function and increased susceptibility to the effects of cigarette smoke. PloS one 2010, 5(1): e8660.

50. Wyman PA, Moynihan J, Eberly S, Cox C, Cross W, Jin X, et al. Association of family stress with natural killer cell activity and the frequency of illnesses in children. Archives of pediatrics & adolescent medicine 2007, 161(3): 228–234.

51. Riaz N, Havel JJ, Makarov V, Desrichard A, Urba WJ, Sims JS, et al. Tumor and Microenvironment Evolution during Immunotherapy with Nivolumab. Cell 2017, 171(4): 934–949 e916.

52. Kim ST, Cristescu R, Bass AJ, Kim KM, Odegaard JI, Kim K, et al. Comprehensive molecular characterization of clinical responses to PD-1 inhibition in metastatic gastric cancer. Nat Med 2018, 24(9): 1449-+.

53. Chen DS, Mellman I. Elements of cancer immunity and the cancer-immune set point. Nature 2017, 541(7637): 321–330.

54. Wargo JA, Reddy SM, Reuben A, Sharma P. Monitoring immune responses in the tumor microenvironment. Current opinion in immunology 2016, 41: 23–31.

55. Dunn GP, Dunn IF, Curry WT. Focus on TILs: Prognostic significance of tumor infiltrating lymphocytes in human glioma. Cancer immunity 2007, 7: 12.

56. Eerola AK, Soini Y, Paakko P. A high number of tumor-infiltrating lymphocytes are associated with a small tumor size, low tumor stage, and a favorable prognosis in operated small cell lung carcinoma. Clin Cancer Res 2000, 6(5): 1875–1881.

57. Vesely MD, Kershaw MH, Schreiber RD, Smyth MJ. Natural innate and adaptive immunity to cancer. Annual review of immunology 2011, 29: 235–271.

58. Crispo A, Brennan P, Jockel KH, Schaffrath-Rosario A, Wichmann HE, Nyberg F, et al. The cumulative risk of lung cancer among current, ex- and never-smokers in European men. British journal of cancer 2004, 91(7): 1280–1286.

59. Brennan P, Crispo A, Zaridze D, Szeszenia-Dabrowska N, Rudnai P, Lissowska J, et al. High cumulative risk of lung cancer death among smokers and nonsmokers in Central and Eastern Europe. American journal of epidemiology 2006, 164(12): 1233–1241.

60. Bjorkstrom NK, Ljunggren HG, Michaelsson J. Emerging insights into natural killer cells in human peripheral tissues. Nature reviews Immunology 2016, 16(5): 310–320.

61. Rock KL, Reits E, Neefjes J. Present Yourself! By MHC Class I and MHC Class II Molecules. Trends in immunology 2016, 37(11): 724–737.

62. Pan D, Kobayashi A, Jiang P, de Andrade LF, Tay RE, Luoma AM, et al. A major chromatin regulator determines resistance of tumor cells to T cell-mediated killing. Science 2018, 359(6377): 770-+.

63. Brock G, Datta S, Pihur V, Datta S. clValid: An R package for cluster validation. J Stat Softw 2008, 25(4): 1–22.

